# Low level of viral intra-host diversity is associated with severe SFTS disease

**DOI:** 10.1101/2025.04.28.650738

**Authors:** Xuemin Wei, Hongfeng Li, Lintao Sai, Li Song, Xinyi Gao, Chao Li, Xinyu Zhang, Yuanyuan Shen, Shuo Li, Li Liu, Tanya Golubchik, Yifei Xu

**Affiliations:** Department of Microbiology, School of Public Health, Cheeloo College of Medicine, Shandong University, Jinan, Shandong, China; Department of Infectious Diseases, Qilu Hospital of Shandong University, Jinan, Shandong, China; Suzhou Research Institute of Shandong University, Suzhou, Jiangsu, China; Big Data Center, Nanfang Hospital, Southern Medical University, Guangzhou, Guangdong, China; Sydney Infectious Diseases Institute, School of Medical Sciences, Faculty of Medicine and Health, University of Sydney, Sydney, Australia; Big Data Institute, Nuffield Department of Medicine, University of Oxford, Oxford, UK

**Author notes:** Contributed equally (Xuemin Wei, Hongfeng Li, Lintao Sai).

**Keywords:** intra-host diversity, iSNV, SFTSV, disease severity, selection pressure

## Abstract

Severe fever with thrombocytopenia syndrome virus (SFTSV) is an emerging tick-borne virus that can cause severe clinical symptoms and high fatality. However, the dynamics of SFTSV intra-host diversity during infection and its association with disease severity remain unexplored. Here, we characterized SFTSV iSNVs in longitudinal clinical samples from patients with distinct clinical outcomes. Our analyses revealed that SFTSV intra-host diversity was associated with disease severity and showed distinct patterns in severe and mild cases. Severe cases exhibited paradoxically lower SFTSV intra-host diversity despite higher viral loads, likely driven by reduced selection pressure. Notably, iSNVs linked to viral replication emerged in severe cases, while iSNVs associated with immunity invasion arose in mild cases. Machine learning models showed that intra-host diversity can contribute to classification of SFTS by severity. Our findings, through the lens of within-host viral populations, highlight variations in viral-host interactions during SFTSV infection and provide potential biomarkers for predicting SFTS severity.

## Introduction

Severe fever with thrombocytopenia syndrome (SFTS) is an emerging tick-borne infectious disease caused by SFTSV, a segmented and negative-strand RNA virus belonging to the genus *Bandavirus* within the family *Phenuiviridae*. SFTS was first identified in China in 2009^1^ and has since affected a rapidly growing number of people, with an expanding geographic distribution, including Japan, Korea, Vietnam, Thailand, and Myanmar^2–4^. SFTS can cause severe clinical symptoms, including hemorrhagic fever, encephalitis, and multiple organ failure, with a high case-fatality rate of 12-50%^5,6^.

Tick-to-human transmission has been recognized as the primary infection route for SFTSV, with *Haemaphysalis longicornis* identified as the main tick vector^7^. Given the risk of epidemic spread, the World Health Organization included SFTSV in a list of priority pathogens with the potential to trigger an international public health emergency in both 2017 and 2024^8^. At present, there is no specific antiviral therapy or approved vaccine available for the treatment or prevention of SFTS. The viral-host interactions and pathogenesis mechanisms of SFTSV remain poorly understood.

Understanding the pathogenesis of SFTSV requires studies of the evolutionary capacity of the virus, which is a combination of genetic variation produced during an individual infection (intra-host single-nucleotide variants, iSNVs) and the propensity of new variants to spread through a population. Most SFTSV studies have been conducted using consensus sequences, focusing on population-level genetic diversity^9–11^. Viral diversity within individual hosts, though less studied, can provide valuable and complementary insight into the generation and spread of novel viral variants, high resolution inference of transmission linkages, drug resistance, and pathogenesis^12–19^. Little is known about the patterns of SFTSV diversity during the course of infection, or the relationship between intra-host variation and severity of SFTS disease. Addressing this gap is a priority for understanding the pathogenesis and developing effective anti-SFTS therapeutic strategies.

In this study, we looked beneath the consensus genomic sequence to characterize SFTSV within-host diversity in longitudinal clinical samples from 73 patients with distinct clinical outcomes, including 40 with mild and 33 with severe disease. Using deep sequencing, we compared the pattern of intra-host diversity within and between patients with different disease severity, to explore the evolutionary capacity of SFTSV and the relationship between viral diversity and disease severity. Our work expands our knowledge of the SFTSV viral-host interaction and provides new insights into pathogenesis.

## Methods

### Patient information and sample collection

Serum samples were collected from 73 SFTS patients in Jinan, Shandong Province, China between March 2022 and August 2024. SFTS infection was confirmed by RT-PCR testing of serum samples. All patients reported a history of tick bite or contact with infected animals within two weeks prior to disease onset. During post-diagnosis, patients received treatment and management in the ward, with standardized electronic medical records used to collect demographic and hospital laboratory data. This study was approved by the ethical committees of Qilu Hospital and Shandong Province Public Health Clinical Center.

Patients were classified into mild and severe groups based on illness severity. Severe cases were defined by dysfunction in three or more of the following organ systems^20,21^: (1) secondary pulmonary infection requiring high-flow oxygen therapy or ventilator-assisted ventilation; (2) secondary bloodstream infection leading to septic shock necessitating vasopressor therapy; (3) hepatic dysfunction, indicated by serum bilirubin ≥ 2-3 mg/dL or liver function tests ≥ twice the normal range; (4) acute kidney injury, defined as oliguria ≤ 400 mL/24 hours or increased creatinine levels (≥ 2-3 mg/dL); (5) hemorrhagic tendency, characterized by an activated partial thromboplastin time (APTT) increase > 25% or platelet counts ≤ 30×10^9^ /L; (6) neuropsychiatric symptoms; and (7) heart failure.

### Viral load measurements and genomic sequencing

Viral RNA was extracted using the QIAamp Viral RNA Mini Kit (Qiagen, Hilden, Germany) according to the manufacturer’s instructions. Real-time RT-qPCR was performed to target the highly conserved region of SFTSV (DaAn Gene Co., Guangzhou, China). A cycle threshold (Ct) value less than 35 indicated a positive test for SFTSV. Positive samples were sequenced, and viral RNA was extracted again using the same protocol. Sequencing libraries were constructed using the IGT® Fast Stranded RNA Library Prep Kit v2.0 (iGeneTech, Beijing, China). Briefly, RNA was fragmented and reverse-transcribed into first-strand DNA using random primers, followed by second-strand DNA synthesis. The resulting double-stranded DNA was end-repaired, A-tailed at the 3’ ends, and ligated to adapters. After PCR amplification for 15 cycles with unique dual indexes, the PCR products were purified using IGT® Pure Beads (iGeneTech, Beijing, China). The purified libraries were further enriched using biotinylated probes targeting the whole viral genome with the TargetSeq One® Hyb & Wash Kit v2.0 (iGeneTech, Beijing, China) according to the manufacturer’s instructions. The final viral-enriched libraries were sequenced on an Illumina NovaSeq 6000 platform to generate 2 × 150 bp paired-end reads.

### Calling of iSNVs

Paired-end reads were mapped to the reference genome (GenBank accession: HM745930, HM745931, and HM745932) using BWA mem (v0.7.16) with default parameters^22^. Duplicated reads were marked using Picard MarkDuplicates (v2.10.10, http://broadinstitute.github.io/picard) with default settings. The base composition of each position was summarized from the mapped reads using pysamstats (v1.1.2, https://github.com/alimanfoo/pysamstats). Samples with sequencing depth greater than 100 and reference coverage greater than 10% were used to identify iSNVs. Minor allele frequency (MAF) was defined as one minus major allele frequency. The mean MAF per sample was determined by dividing the total MAF across all genomic positions by the total number of positions. High-quality iSNVs were ensured through the application of several criteria: (1) base quality greater than 20; (2) sequencing coverage of paired-end mapped reads greater than 100; (3) at least 15 reads support the minor allele; (4) minor allele frequency greater than 5%; and (5) strand bias ratio of reads with the minor allele and reads with major allele less than ten-fold. To minimize errors, serial adjacent iSNVs (distance less than 50 bp) containing more than five iSNVs were excluded.

Consensus sequences were generated with a minimum of 10-fold mapping coverage and supported by at least 50% of reads at a given position. The sequences obtained in this study, together with representative sequences from GenBank, were aligned using MAFFT v7.310^23^. A maximum likelihood phylogenetic tree was constructed using IQ-TREE v2.0.3 with the best-fit substitution model determined by the software^24^. Branch support was assessed with ultrafast bootstrapping.

### Validation of iSNVs by re-sequencing

To assess the reproducibility of the iSNVs, two independent libraries were prepared for 28 samples. The consistency of iSNVs detected in the two sequencing experiments was evaluated using a scatter plot. iSNVs with a MAF below 5% were generally indistinguishable from background noise, whereas those with a MAF above 5% showed high concordance between the two experiments (Fig. S1a-b).

### Evaluating iSNV genomic distance distributions

iSNV positions were simulated in R using the runif function with a uniform distribution based on the number of iSNVs. Both actual and simulated iSNVs were ranked, and genomic distances between neighboring mutant pairs were calculated. The distances between simulated iSNVs followed a Poisson distribution. To evaluate the distribution of distances between adjacent iSNVs, the Kolmogorov-Smirnov test was used to compare the observed distribution with the expected distribution under a Poisson model. The iSNV density in a given region was calculated by dividing the total number of identified iSNVs by the number of samples, and then further dividing by the length of the region.

### Analysis of intra-host diversity

SFTSV intra-host diversity was estimated using pairwise nucleotide diversity (*π*)^25,26^. The proportion of pairwise nucleotide differences (*D*) at each genomic position was calculated as:

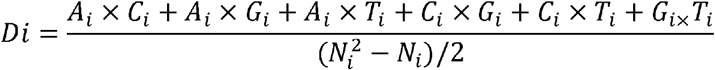

where *A*_*i*_,*C*_*i*_, *G*_*i*_, and *T*_*i*_ represent the copy number of allele A, C, G, and T, respectively, and *N*_i_ is the total count of the four alleles (i.e. depth of coverage) at a given locus i, so *N*_*i*_ = *A*_*i*_ +*C*_*i*_+ *G*_*i*_ +*T*_*i*_. Loci with a total count of less than 100 were excluded. Pairwise nucleotide diversity across a genome (π) was then calculated as

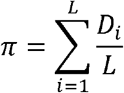

where L is the number of genomic positions with a read depth over 100×.

### Selection pressure and adaptive mutation

Ka and Ks were determined using KaKs Calculator (v2.0.1) with the MYN model^27^, which accounted for the rates of transitions and transversions as well as codon frequency bias. Candidate functional iSNVs were identified by compiling mutations reported in literature to affect SFTSV function. These included mutations linked to viral replication, immune evasion, and other functional sites. The mutations were summarized in Table S1.

### Statistical analysis

Comparisons of continuous variables between groups were conducted using two-tailed Mann-Whitney *U*-tests (two groups) or Kruskal-Wallis tests (three groups). The relationship between two variables was assessed using Pearson correlation analysis. Linear regression models were constructed to evaluate the relative influence of Ct value, genotype (C1, C2, and C3; Fig. S3), days post symptom onset, and disease severity on intra-host diversity, with adjustments for age and gender (significance threshold: *P* < 0.05). Intra-host diversity was plotted against Ct value and a linear model was used to adjust for Ct values. Additionally, logistic regression models were used to assess the effect of viral and clinical parameters on disease severity, with age and gender as covariates (*P* < 0.05/34 = 0.0015). Temporal changes of variables were assessed through ordinary least-squares linear regression. All statistical analyses were performed using R (v4.3.1).

### Machine learning

Logistic regression and random forest models were constructed using viral parameters and clinical parameters to distinguish severe and mild SFTS cases. Prior to modeling, variables with over 50% missing values were excluded, while remaining variables were interpolated using the random forest approach with the *missForest* package. Highly correlated variables (i.e., Pearson’s *r* > 0.7) were removed (Fig. S2), and numeric variables were centered. The dataset was randomly divided into training (60%) and test set (40%) based on disease severity. Least Absolute Shrinkage and Selection Operator (LASSO) regression was applied to identify the most predictive variables through analysis of multiple random training subsets with varying shrinkage parameters. Four-fold stratified cross-validation was performed, and the average area under the receiver operating characteristic curve (AUC) was plotted against the number of model variables. The optimal model was selected based on the elbow point in the plot, indicating peak performance without overfitting^28^. Additionally, all-possible regressions were applied to identify the best model among top-ranked variables. The modeling process was replicated using the random forest algorithm. Furthermore, the AUC of the best model with and without viral biomarkers was statistically compared using the two-sided DeLong test^29^.

## Results

### Most SFTSV infections have low intra-host diversity

We sequenced 126 serum samples collected from 73 SFTS patients from 2022 to 2024. After removing three samples with low viral genomic coverage or abnormal number of iSNVs, 123 samples were retained in the analyses. These samples represented two groups: 1) 51 samples collected from 51 patients, including 30 mild cases and 21 severe cases; 2) 72 longitudinal samples from 21 patients, including 9 mild cases and 12 severe cases (Fig. 1a). Each patient was sampled at least two times (mean 3.43, range 2-5), with sampling intervals ranging from one to six days.

**Figure 1.**
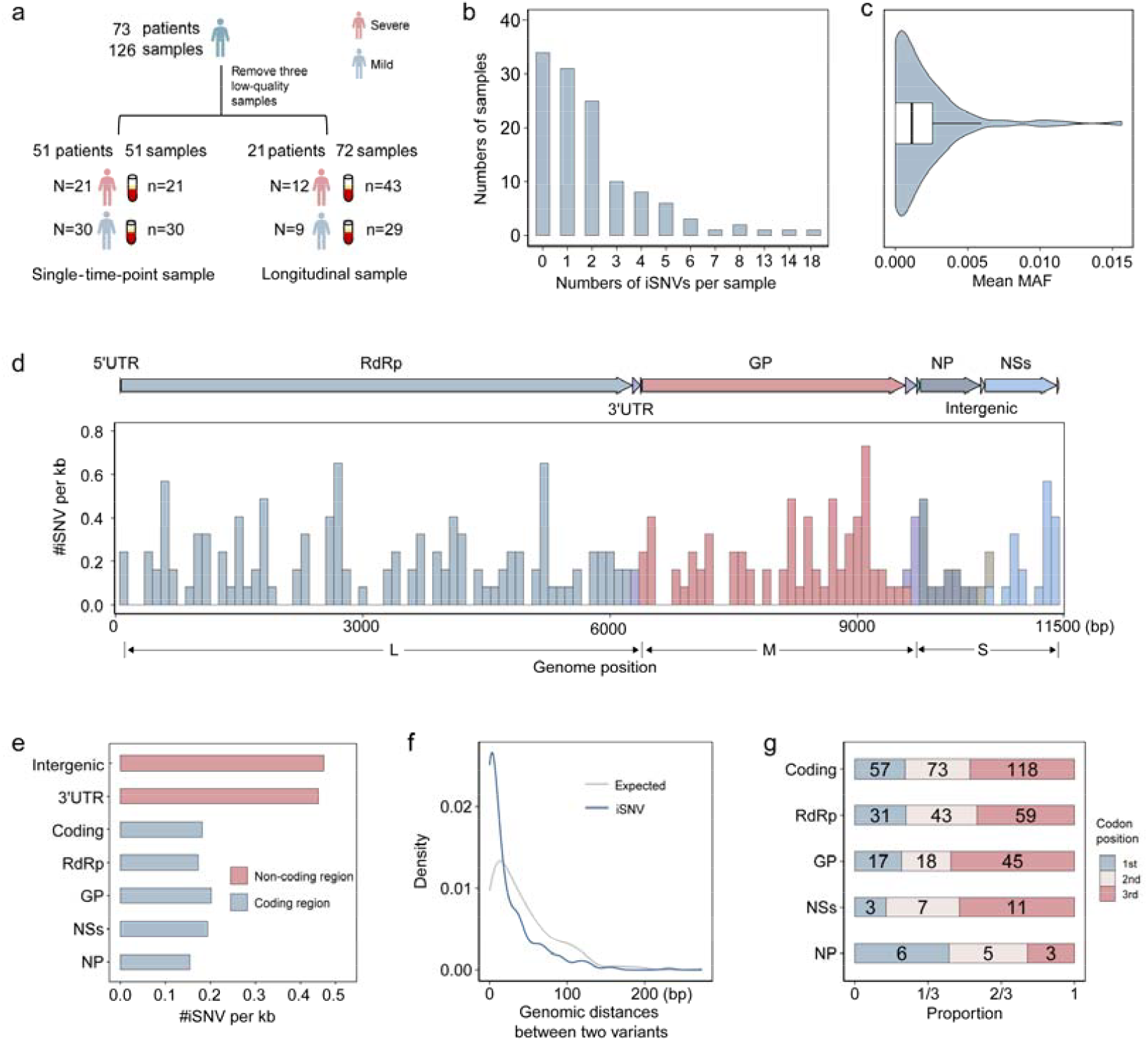
Distribution of identified iSNVs in SFTSV. a) Flowchart of sample collection and sequencing. b) Distribution of iSNVs number in the samples. c) Distribution of mean MAF in the samples. d) Distribution of iSNVs density along the genome, counted using a window of 100 bp. e) Normalized number of iSNVs in coding and non-coding regions. f) Distribution of genomic distance between two variants in expected and observed iSNVs. The blue line represents the observed iSNVs, while the gray line shows the expected distribution modeled by a Poisson process. g) Proportion of iSNVs at different codon positions in each gene. The number of iSNVs in each category is marked on the corresponding bar.

The mean sequencing depth of each sample showed a negative correlation with Ct values (R-square [R^2^] = 0.59, *P* < 0.001; Fig. S1c). The number of iSNVs in each sample was not affected by genomic coverage (R^2^ < 0.001, *P* = 0.93; Fig. S1d). While a statistical correlation was observed between the number of iSNVs and mean sequencing depth, the negative correlation was weak (R^2^ = 0.137, *P* < 0.001; Fig. S1e).

We identified a total of 259 iSNVs from 123 samples, with a median of one iSNV per sample (range 0-18; Fig. 1b). 34 samples had no iSNVs. Mean MAF per sample ranged from 0-0.0156% (median 0.0011%, Fig. 1c). We examined the distribution of iSNVs across the SFTSV genome and found an overall density of 0.19 iSNVs/kb (Fig. 1d). When normalized by length, the intergenic region showed the highest iSNV density (0.45 iSNVs/kb), followed by 3’-UTR (0.44 iSNVs/kb) and coding region (0.18 iSNVs/kb; Fig. 1e). The majority (248, 95.75%) of iSNVs was located in coding regions, which cover 96.58% of the whole genome. Within the coding regions, the glycoprotein (GP) gene displayed a higher iSNV density (0.20 iSNVs/kb) compared to other genes. This gene encodes a polyprotein precursor that is cleaved into Gn and Gc envelope glycoproteins, which is crucial for viral entry and immune evasion.

We calculated the genomic distances between allele pairs of iSNVs to investigate whether their distribution across the genome was non-random. The fitted density curve showed a significant difference from randomly generated mutations (Kolmogorov-Smirnov test, *P* < 0.001; Fig. 1f), indicating that distribution of iSNVs may be driven by positive selection or mutational hotspots. We also analyzed the distribution of iSNVs at distinct codon positions, identifying 57, 73, and 118 iSNVs at the first, second, and third codon positions, respectively (Fig. 1g). The GP and RNA-dependent RNA polymerase (RdRp) genes had a significantly greater number of iSNVs at the third codon position compared to other positions (Fisher’s exact tests, all *P* < 0.05). Moreover, the MAF of the first and third codons in RdRp was higher than that of the second codon (Mann-Whitney *U*-tests, *P* < 0.001).

iSNVs generally did not occur at the same genomic position in different samples. Among all identified iSNVs, 156 (60.23%) were present in one sample, 26 (10.04%) were shared by two samples, and the remaining 26 (10.04%) were shared by three or more samples (excluding 51 iSNVs shared only by sequential samples from the same patient). We found one iSNV shared by six samples from five patients, with a MAF between 5% and 15%. This iSNV had a A>G substitution at position 2750 of the M segment, causing an amino acid alteration from glutamic acid to glycine at position 911 (E911G) in the GP gene. Phylogenetic analysis showed that our SFTSV consensus sequences were classified into clade C1 (n=35), C2 (n=25), and C3 (n=63). The three segments from the same samples exhibited identical clade assignments (Fig. S3).

### Severe cases have lower intra-host SFTSV diversity independent of viral load

We investigated the relationship between intra-host diversity, Ct value, genotype, days post symptom onset, and disease severity. Intra-host diversity was estimated using pairwise nucleotide diversity (pi), yielding a median of 0.0137 (range 0.0047-0.0260) for the whole dataset. We found that genotype, days post symptom onset, gender, and age were not significantly associated with intra-host diversity (linear regression, *P* > 0.05; Fig. 2a and Fig. S4c).

**Figure 2.**
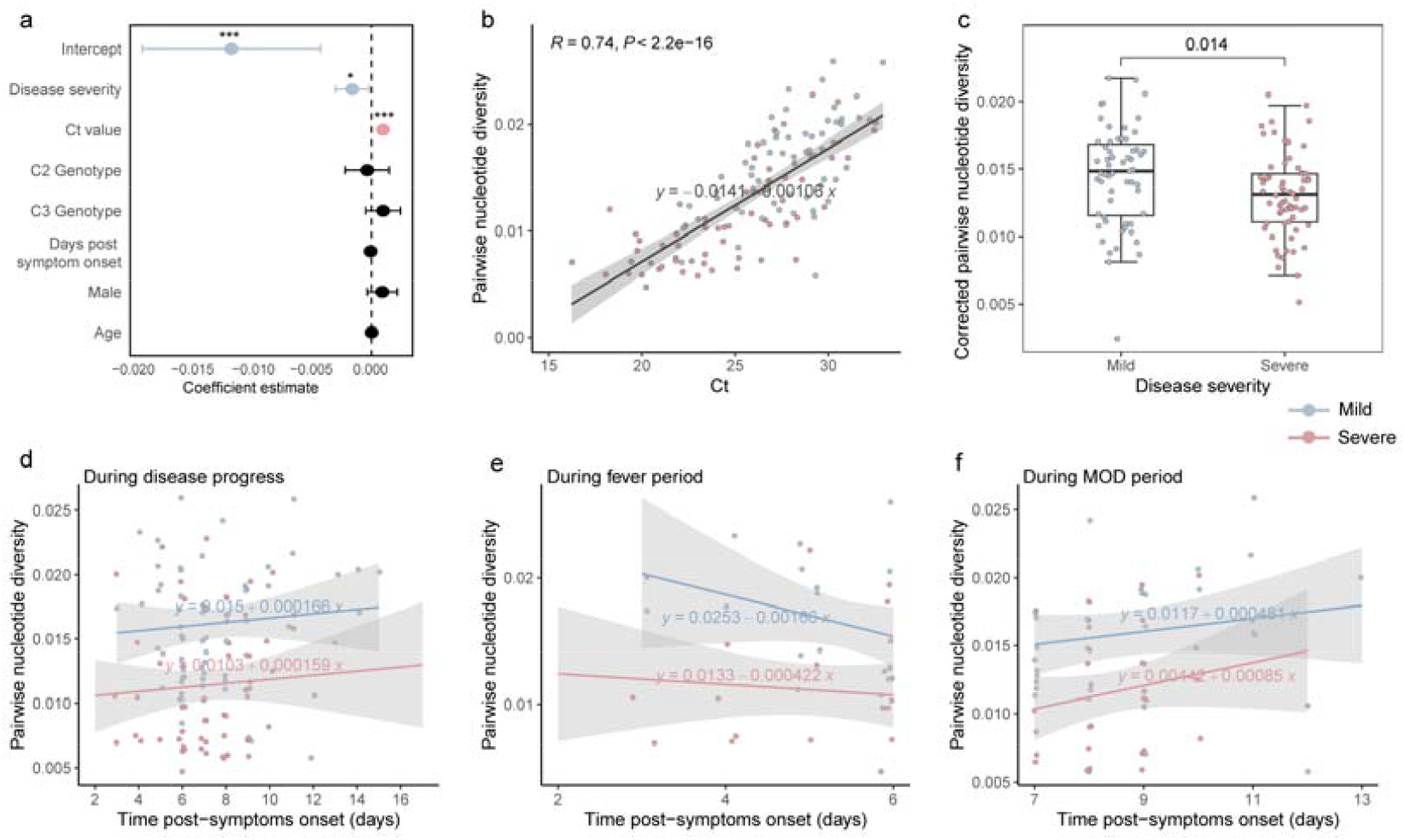
Factors that affect SFTSV intra-host diversity. a) Coefficients and corresponding *P*-values were estimated using the least-squares linear regression. Circles represent the estimates and lines show 95% confidence intervals. \**P*<0.05, \*\**P*<0.01, and \*\*\**P*<0.001. b) Pairwise nucleotide diversity against Ct value. c) Comparison of corrected pairwise nucleotide diversity between mild and severe groups. Pairwise nucleotide diversity against time after symptom onset during d) disease process, e) fever period, and f) MOD period. In b)-f) blue indicates mild group and red indicates severe group.

Intra-host diversity was positively correlated with Ct value of the samples, that is, negatively correlated with viral load (Pearson correlation, *r*=0.742, *P* < 0.001; linear regression, β = 9.54e-4, *P* < 0.001; Fig. 2a and Fig. 2b). Samples from severe cases had higher viral loads than those from mild cases. Notably, our analysis revealed that severe group exhibited significantly lower intra-host diversity compared to mild group (Mann-Whitney *U*-tests, *P* < 0.001; linear regression, β = −0.0016, *P* = 0.026; Fig. 2a and Fig. S4a). This difference remained significant after adjusting for the Ct effects (Mann-Whitney *U*-tests, *P* = 0.014; linear regression, β = −0.0016, *P* = 0.024; Fig. 2c, Fig. S4b and Fig. S4c).

### Dynamics of intra-host diversity during distinct stages of infection

The longitudinal samples provided opportunity to track changes in intra-host diversity over the course of SFTSV infection. Putting all samples together, we observed that intra-host diversity accumulated during infection, rising from 0.012 (mean pairwise nucleotide diversity) on the first day to 0.014 ten days after symptom onset (Fig. S4d). Notably, severe cases exhibited lower intra-host diversity on the initial day of symptom onset, and a similar accumulation rate compared to mild cases (Fig. 2d). We then analyzed the changes during the fever period and multiple organ dysfunction (MOD) period^30^. Intra-host diversity declined during the fever period in both mild and severe groups (Fig. 2e), but accumulated during MOD period, with a faster accumulation rate in severe group (Fig. 2f). We observed a similar pattern when only considering the temporal samples from 21 participants who had serial samples collected. We also evaluated each set of serial samples collected from the same patient. Intra-host diversity showed a consistent increase in approximately half of the patients (mild/severe: 6/4), whereas it exhibited random variations and fluctuations in the other half (Fig. S5).

### Functional iSNVs

By analyzing mutations reported in the literature to affect SFTSV function, we identified a total of seven potentially functional candidate iSNVs that exhibited a pattern specific to disease severity in our dataset (Fig. 3a).

**Figure 3.**
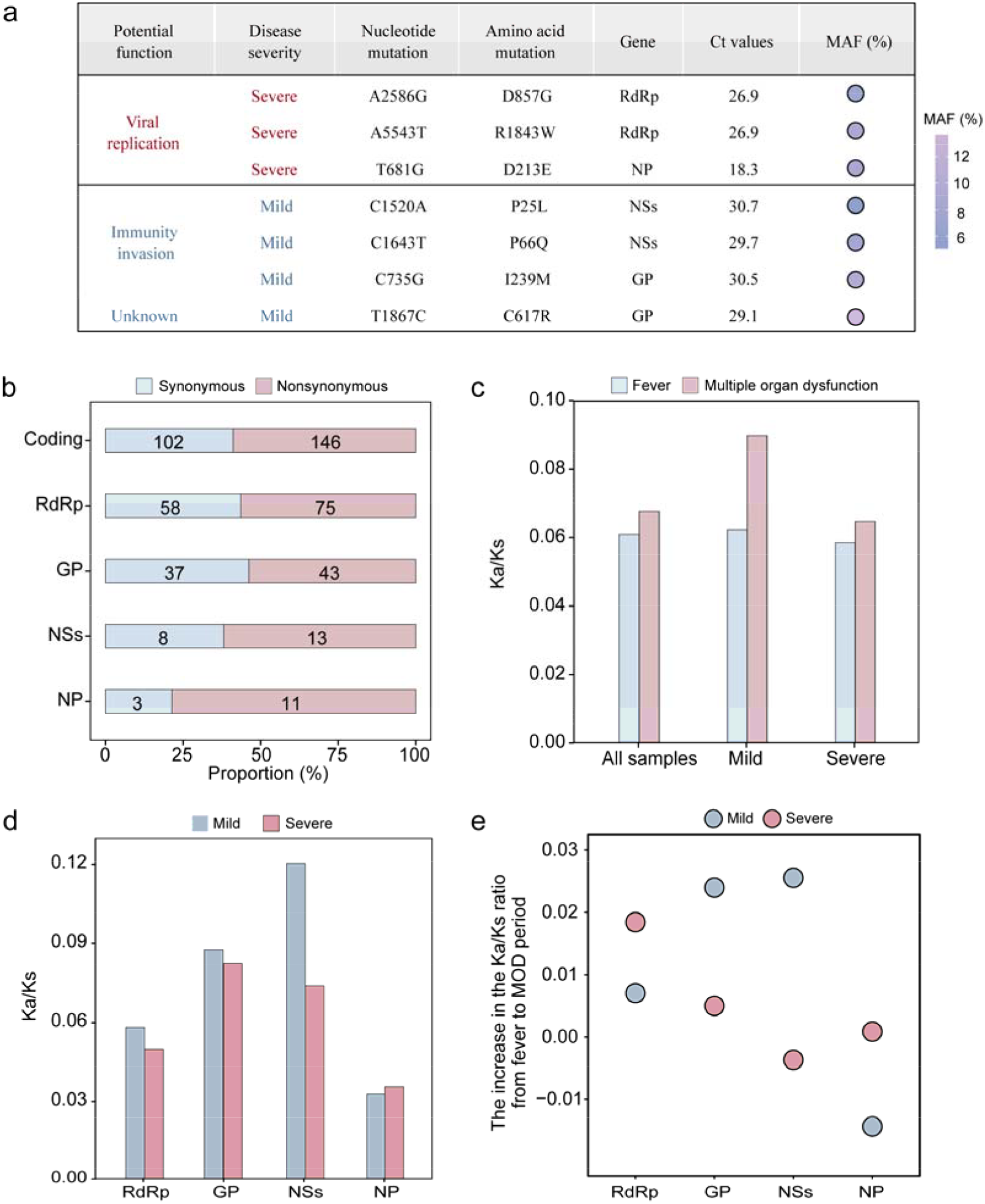
Selection pressures for iSNVs. a) Identification of potential functional iSNVs in mild and severe cases. b) Proportion of nonsynonymous and synonymous iSNVs for each gene. The number of iSNVs in each category is marked on the corresponding bar. c) Ka/Ks ratios for all samples, mild group, and severe group during fever and MOD periods. d) Ka/Ks ratio for each gene. e) The increase in the Ka/Ks ratio from the fever to the MOD period in mild and severe groups.

- The severe cases comprised three functional iSNVs that were linked to viral replication. The D857G mutation (nucleotide mutation: A2586G) in RdRp gene and D213E mutation (T681G) in nucleoprotein (NP) gene, both located at N6-methyladenosine (m^6^A) modification sites, can enhance SFTSV replication^31^. The R1843W mutation (A5543T), located in the RdRp cap-binding pocket, was hypothesized to decrease RNA replication and transcription^32^.
- The mild cases comprised four functional iSNVs that were potentially associated with viral immune evasion. The P66Q (C1643T) and P25L (C1520A) mutations in the nonstructural proteins (NSs) gene contribute to immune evasion by suppressing IFN-β promoter activity^33,34^. The I239M (C735G) mutation, situated within an antigenic epitope of the GP gene, could facilitate immune evasion^35^. The remaining C617R (T1867C) mutation in the GP gene could affect Gc protein expression, although its exact role in viral immune evasion remains uncertain^36^.

These results suggested that the within-host populations of SFTSV may be driven by distinct selective pressures in severe and mild cases.

### Selection pressure for iSNVs

We next analyzed changes in nonsynonymous and synonymous mutations during infection. Nonsynonymous iSNVs accumulated faster than synonymous iSNVs in both severe and mild groups (Fig. S6a). The accumulation rate of nonsynonymous iSNVs was lower in severe group than in mild group during the fever and MOD periods. For all samples, we identified 146 nonsynonymous iSNVs and 102 synonymous iSNVs, with an overall ratio of 1.4 (Fig. 3b). This ratio varied across genes, with the NSs gene showing the greatest ratio (4.3), significantly surpassing other parts in the genome (Fisher’s exact test, *P* = 0.001). The ratio of nonsynonymous to synonymous iSNVs was slightly lower in severe group (1.2) compared to the mild group (1.4; Fig. S6b).

We also measured the ratio of Ka to Ks, which compares the rate of nonsynonymous substitutions per nonsynonymous site (Ka) with the rate of synonymous substitutions per synonymous site (Ks). The mean Ka/Ks ratios were below one, showing that SFTSV was under purifying selection. The ratios during the MOD period were higher than those during the fever period in both severe and mild groups (Fig. 3c). Notably, severe cases exhibited a lower mean Ka/Ks ratio compared to mild cases. These results suggested that selection pressure on SFTSV increased as the disease progressed, and severe cases experienced lower selection pressure than mild cases.

GP gene showed the greatest Ka/Ks ratio in severe group, whereas NSs gene exhibited the greatest ratio in mild group (Fig. 3d). When comparing the mild and severe groups, we found that the NSs gene displayed the most noticeable difference in mean Ka/Ks ratios. As SFTS progressed from fever to MOD period, the NSs gene showed the greatest increase in Ka/Ks ratio in mild group (Fig. 3e). In contrast, the RdRp gene exhibited the greatest increase in Ka/Ks ratio in the severe group.

We analyzed the mutational types of iSNVs and found that top four iSNVs were transitions (A<>G and C<>T), which together accounted for 79.15% (205/259) of total iSNVs (Fig. 4a). Transition mutations exhibited significantly higher iSNVs/kb compared to transversions (Mann-Whitney U test, *P* < 0.001; Fig. 4c), whereas no significant difference in MAF was observed (*P* = 0.15; Fig. 4d). The overall transition/transversion ratio (*ti/tv*) was 3.8 (205:54) for all samples, consistent with short evolutionary timescale^37^. Additionally, nonsynonymous substitutions were more enriched in transitions than in transversions (Chi-Squared test, *P* = 0.002).

**Figure 4.**
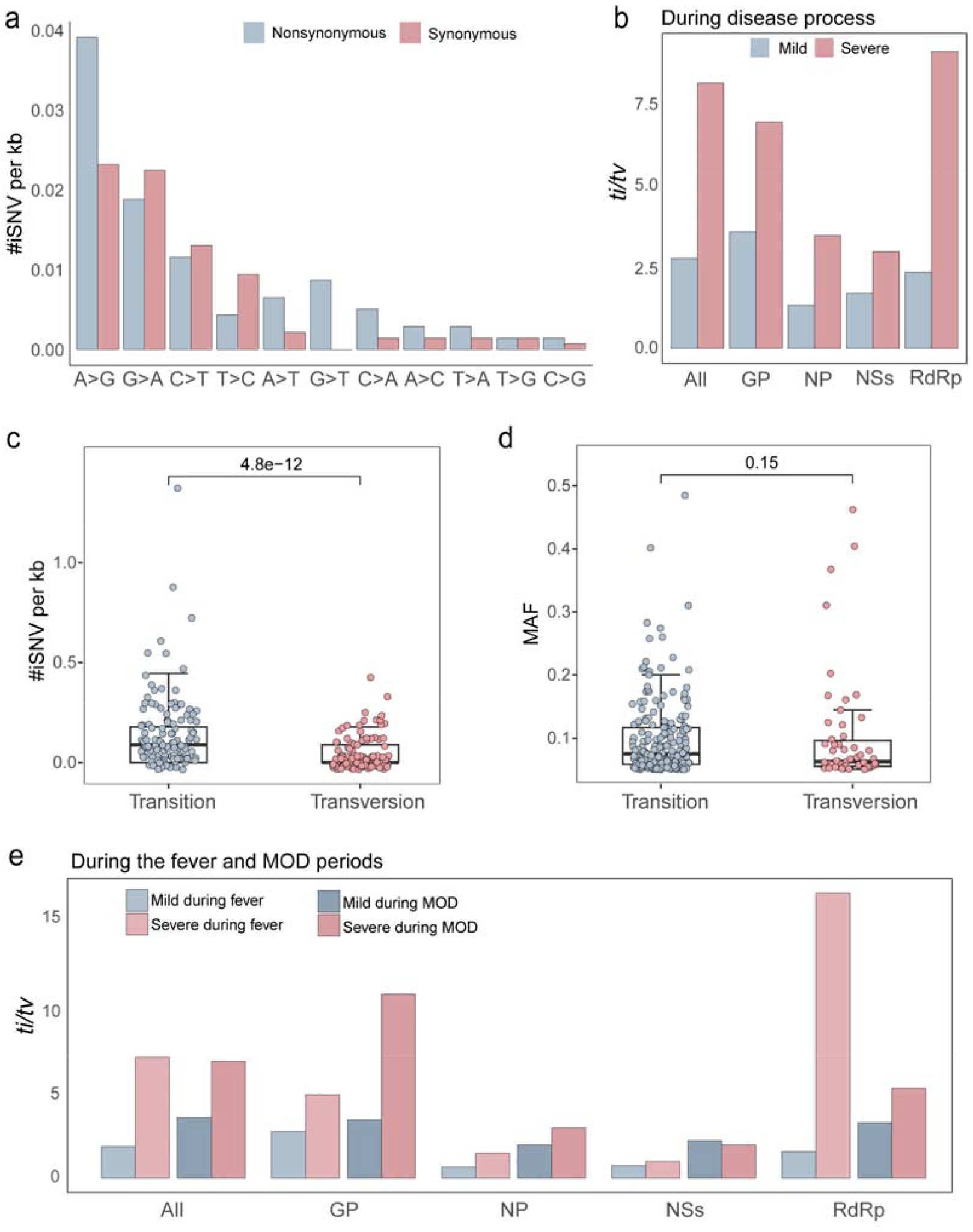
Transition/transversion ratio for SFTSV. a) Number of iSNVs for different mutation types. b) The *ti/tv* of different genes in the mild and severe groups during disease process. Comparison of c) iSNV/kb and d) MAF between transition and transversion. e) The *ti/tv* in mild and severe groups during the fever and MOD periods.

Comparison showed that the severe group exhibited higher *ti/tv* compared to mild group during both fever and MOD periods, except for the NSs gene during the MOD period (Fig. 4b and Fig. 4e). As SFTS progressed from fever to MOD period, *ti/tv* increased in mild group but slightly decreased in severe group (Fig. 4e). Notably, among all genes, the RdRp gene showed the greatest *ti/tv* increase in mild group but the most decrease in the severe group (Fig. 4e).

### Machine learning models to predict SFTS severity

Finally, we investigated whether intra-host diversity could be used in machine learning modeling for classifying SFTS severity. Based on logistic regression analysis adjusted for age and gender, we identified 14 variables that were significantly associated with disease severity (*P* < 0.05). Three variables were viral parameters, including pairwise nucleotide diversity (β = −244.92, *P* < 0.001), viral load as represented by Ct values (β = −0.33, *P* < 0.001), and iSNVs/kb (β = −5.10, *P* = 0.014). The remaining eleven variables were clinical parameters, such as sialic acid (β = −0.13, *P* = 0.004) and adenosine deaminase (ADA, β = 0.085, *P* = 0.0015; Fig. 5a). We constructed two machine learning models, one with and one without viral parameters, and tested their performance using cross-validation. The best performing models included pairwise nucleotide diversity, age, and sialic acid (Fig. 5b). Notably, the level of intra-host diversity, represented by pairwise nucleotide diversity, had the second-highest feature importance in the model after sialic acid (Fig. 5b). Incorporating viral parameters into either logistic regression or random forest models significantly improved their predictive performance on independent test data (Fig. 5), increasing AUCs for logistic regression to 0.83 versus 0.70 (*P* = 0.029) and random forest to 0.78 versus 0.69 (*P* = 0.027; Fig. 5c and Fig. 5d).

**Figure 5.**
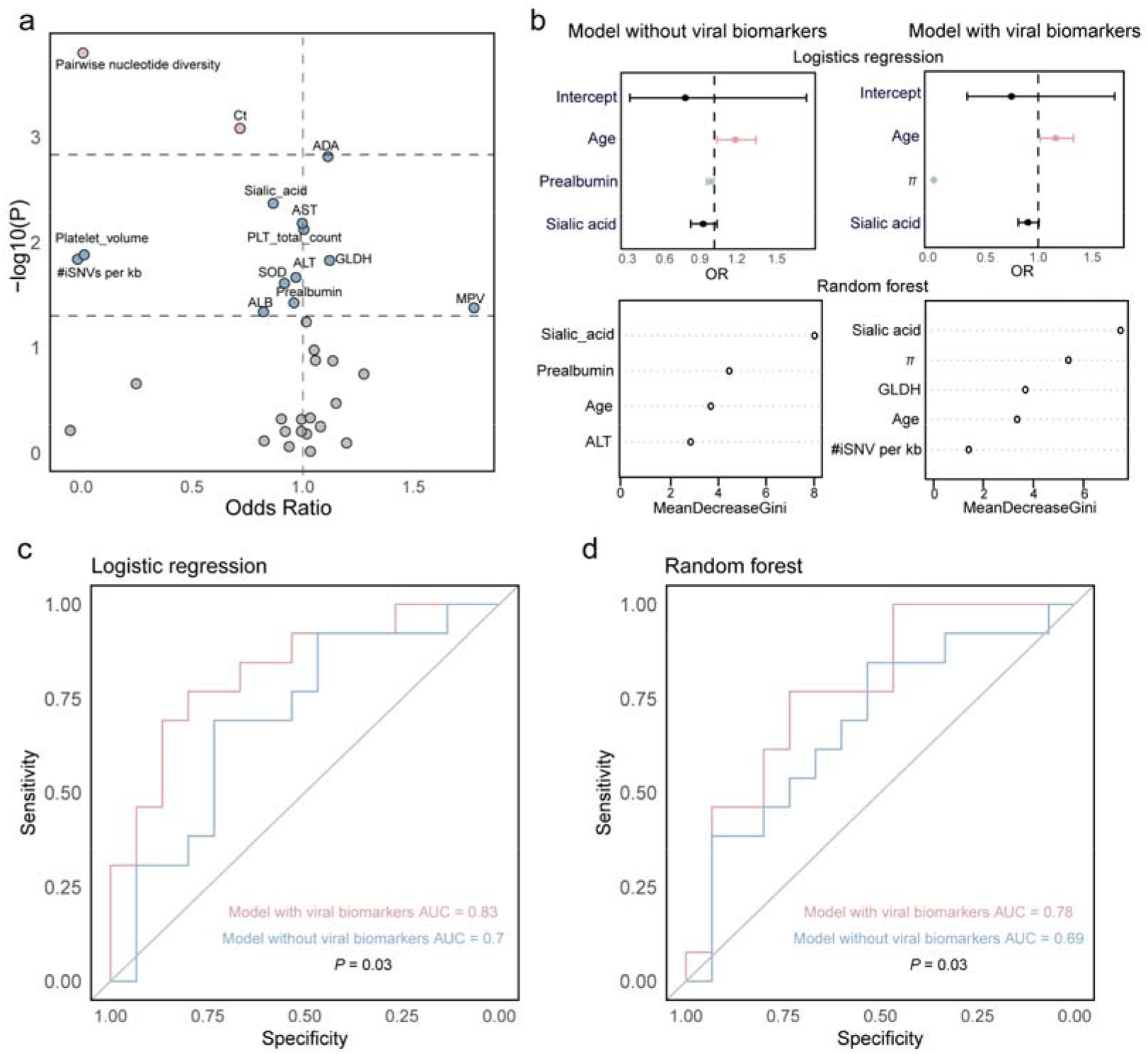
Classification of SFTS by severity using machine learning models. a) Volcano plot of viral and clinical parameters. Red: *P* < 0.05/34 = 0.0015; blue: *P* < 0.05; black: *P* < 0.05. b) Optimal logistic regression and random forest models for predicting the severity of SFTS. Red: a positive correlation with disease severity; blue: a negative correlation with disease severity. π represents pairwise nucleotide diversity. MeanDecreaseGini quantifies the importance of each variable within the random forest model, with higher scores indicating greater contributions to the model’s predictive capability. Predictive performance of c) the best logistics regression models and d) the best random forest models evaluated with and without viral biomarkers.

## Discussion

Intra-host viral diversity offers valuable insights into the emergence and dissemination of novel viral variants, enabling high-resolution inference of transmission networks, understanding of drug resistance, and elucidation of pathogenesis. In this study, we conducted, to the best of our knowledge, the first investigation of SFTSV within-host populations dynamics during infection, and explored the potential association between within-host populations and disease severity.

### Comparison of intra-host diversity between SFTSV and other RNA viruses

We identified a total of 259 iSNVs from 123 SFTSV samples, with a median density of 0.19 iSNVs/kb, which is comparable to, or slightly lower than those previously reported for other RNA viruses, including SARS-CoV-2, influenza A virus, Ebola virus, and Yellow Fever virus^12,38–40^. This finding is consistent with previous research showing that SFTSV exhibits lower evolutionary rate than other RNA viruses at the population level^41^. Like all arthropod-borne viruses (arboviruses), SFTSV is under selective pressure to maintain replication competence in both the vector and host species, which limits the population-scale evolutionary rate.

### Accumulation of intra-host diversity during infection

The extent of intra-host diversity depends on viral load and duration of infection. In line with previous findings for SARS-CoV-2, another acute virus^12^, we found that SFTSV intra-host diversity accumulated during infection and was negatively correlated with viral load. Viral loads typically decrease as infection progresses after the initial peak, whereas observed intra-host diversity may increase, for example in response to ongoing immune evasion, as well as due to a stochastic sampling effect of relatively few template molecules. Degradation and deamination of viral RNA as infection progresses may also contribute to the observed increased diversity in samples with low viral load.

### Weak immune response in severe cases could lead to lower SFTSV intra-host diversity

Intra-host viral diversity reflects the evolutionary capacity of the viral population, can impact its replication capacity, pathogenicity, and immune evasion abilities, ultimately affecting disease severity. Contrary to classic assumptions that higher viral diversity drives pathogenesis, our study revealed an inverse relationship between SFTSV within-host diversity and disease severity, with severe cases exhibited lower intra-host diversity than the mild cases. This finding aligns with patterns reported in human papillomavirus infection, where high-grade cervical lesions harbored fewer iSNVs^42^, and in H7N3 avian influenza, where highly pathogenic strains showed lower within-host diversity compared to low pathogenicity counterparts^43^. Notably, this contrast with poliovirus, where diminished intra-host diversity correlated with attenuated pathogenicity^17^, and HIV, which leverages high intra-host diversity to evade immunity and accelerate disease progression. These discrepancies likely reflect differences in host-virus interaction dynamics across viruses.

To dissect the evolutionary forces underlying this paradox, we compared SFTSV selection pressures between patients with different disease severity. We observed lower mean Ka/Ks ratio and higher *ti/tv* in severe cases compared to mild cases, suggesting that SFTSV underwent lower selection pressure in severe cases. Moreover, adaptive iSNVs exhibited patterns specific to disease severity: mild cases accumulated immunity-associated iSNVs (e.g., P66Q and P25L mutations in the NSs gene, I239M and C617R mutations in the GP gene), suggesting active viral-host interactions, while severe cases harbored replication-related iSNVs (e.g., D857G and R1843W mutations in the RdRp gene, D213E mutation in the NP gene) that may enhance viral propagation. These results indicated diverging evolutionary trajectories of SFTSV in mild and severe cases, which potentially affected by the differential immune status of the host. Previous studies have attributed SFTS severity to immune deficiency and dysregulation, characterized by exacerbated IFN-I response, T cell depletion, impaired B cell maturation, and inhibited antibody secretion in deceased patients^40–43^.

Based on our observations, we propose that host immune status creates distinct selection landscapes shaping SFTSV intra-host evolution. In severe cases, the weakened immune system fails to effectively control SFTSV, causing persistent viral replication and continued increase in viral loads, with no variant emergence, consistent with an absence of selective pressure. Conversely, in mild cases, the robust immune system is more likely to suppress viral replication but also exert selective pressure to promote the emergence of escape mutations, thereby increasing intra-host diversity. Further studies on the dynamics of SFTSV within-host populations across various disease stages and organs, as well as their interactions with the immune response, will enhance our understanding of how intra-host diversity contributes to the severity of disease.

### NSs gene under greatest selection pressure

Our data showed that the NSs gene exhibited the highest ratio of nonsynonymous to synonymous iSNVs among all genes, consistent with population-level analyses^44^. Moreover, the NSs gene displayed the greatest difference in mean Ka/Ks ratios between severe and mild groups. Previous research has demonstrated that NSs protein sequesters multiple antiviral proteins, including TBK1 (TANK binding kinase 1), RIGI, IKBKE/IKKε (inhibitor of nuclear factor kappa B kinase subunit epsilon), IRF3 (interferon regulatory factor 3), and TRIM25 (tripartite motif containing 25), into inclusion bodies to escape the antiviral innate immune response^45,46^. Additionally, NSs induces complete autophagy and traps host antiviral proteins inside the autophagic vesicles for degradation thereby escaping host innate immune responses^47^. Our findings provide additional evidence supporting that NSs plays an important role in mediating SFTSV pathogenesis.

### Limitations

The dynamic interplay between SFTSV intra-host diversity and host immune responses during the course of infection remains unexplored. Longitudinal characterization of both viral intra-host diversity and concurrent immune parameters through serial sampling could elucidate critical viral-host interactions governing disease progression. In addition, due to the limited sample size, our machine learning model for predicting the SFTS severity requires further validation.

## Conclusions

In summary, our findings demonstrate that SFTSV intra-host diversity is associated with disease severity, reflecting variations in viral-host interactions during infection. Our study offers novel insights into the pathogenesis of SFTSV and presents potential biomarkers for predicting disease severity.

## Acknowledgements

This study was supported by the State Key Research Development Program of China (2023YFC2308500), Taishan Scholars Project (tsqn202306003), Natural Science Foundation of Jiangsu Province (BK20220270), and Suzhou Technology Project (2022SS15). TG is supported by an Investigator Grant (GNT2025445) from the National Health and Medical Research Council, Australia (NHMRC).

## Competing interests

None declared.

## Notes

### Competing Interest Statement

The authors have declared no competing interest.

